# The transcription factor OCT6 promotes the dissolution of the naïve pluripotent state by repressing *Nanog* and activating a formative state gene regulatory network

**DOI:** 10.1101/2023.07.13.548918

**Authors:** Ariel Waisman, Federico Sevlever, Denisse Saulnier, Marcos Francia, Antonella Lombardi, Celeste Biani, María Belén Palma, Agustina Scarafía, Joaquín Smucler, Guadalupe Amín, Alejandro La Greca, Lucía Moro, Gustavo Sevlever, Alejandra Guberman, Santiago Miriuka

## Abstract

Animal development relies on complex gene regulatory networks (GRNs) that govern the nearly irreversible changes that occur during cell differentiation. In this work we aimed to determine key transcription factors (TFs) associated with the dissolution of the naïve pluripotent state and the acquisition of a formative identity. We identified OCT6 as one of the earliest TFs induced during the onset of mouse embryonic stem cell (mESCs) differentiation. To investigate its role, we generated an *Oct6* knockout mESC line, which failed to acquire the characteristic cell morphology associated with the formative state. Transcriptome analysis of differentiating cells revealed nearly 300 differentially expressed genes compared to wild-type cells, including pluripotency TFs *Nanog, Klf2, Nr5a2, Prdm14,* and *Esrrb*, that failed to correctly downregulate. Notably, premature expression of OCT6 in naïve cells triggered a rapid morphological transformation mirroring differentiation, accompanied by self-induction of Oct6 and expression of TFs such as *Sox3, Zic2/3, Foxp1*, as well as the formative genes *Dnmt3A* and *FGF5*. Strikingly, the majority of OCT6 expressing cells did not express NANOG. Gene expression and single molecule RNA-FISH analysis confirmed that this regulation was at the transcriptional level. Collectively, our results establish OCT6 as a key TF in the dissolution of the naïve pluripotent state and support a model where *Oct6* and *Nanog* form a double negative feedback loop which could act as a toggle switch important for the transition to the formative state.

**Highlights:**

1. *Oct6* is rapidly induced as mESCs exit ground state pluripotency.
2. Loss of OCT6 negatively affects the transition to formative pluripotency.
3. Premature expression of OCT6 in mESCs is sufficient to induce a formative-like phenotype.
4. OCT6 and NANOG repress each other forming a double negative feedback loop.

## Introduction

During early mammalian development, cells of the pre-implantation epiblast have the capacity to differentiate into all the cell types of the organism, a property known as pluripotency. This feature persists in the post-implantation epiblast and is present until the time of gastrulation, when the three embryonic germ layers emerge (Osorno et al., 2012). Pluripotency exists in a continuum of states but has recently been subdivided into three subtypes that are linked to the different stages of epiblast development. Thus, pre-implantation epiblast cells are referred to be in a naïve pluripotent state, while early and late post-implantation epiblast cells display formative and primed pluripotency, respectively (Smith, 2017). These different subtypes are characterized by changes in their transcriptional, epigenetic, metabolic as well as functional properties (Morgani et al., 2017).

In mice, the transition between the naïve and formative states can be faithfully recapitulated in vitro with the use of mouse embryonic stem cells (mESCs). They represent an invaluable tool for investigating the mechanisms underlying early mammalian development, as they possess the capacity to self-renew indefinitely while retaining naïve pluripotency (Nichols and Smith, 2009). Importantly, they can be induced to differentiate to epiblast-like cells (EpiLCs) that are in a formative state of pluripotency, similar to the early post-implantation epiblast (Hayashi et al., 2011). The conversion from mESCs to EpiLCs thus constitutes an excellent model for studying the molecular mechanisms of pluripotency conversion and changes in cell identity.

Central to these developmental transitions are the gene regulatory networks (GRNs) that orchestrate the precise control of gene expression, guiding cellular fate decisions. GRNs comprise a complex interplay of transcription factors (TFs), signaling molecules, and epigenetic modifiers that act in concert to regulate the dynamic patterns of gene expression during development (Davidson and Erwin, 2006; Davidson and Levine, 2008). An important aspect of GRNs are the intricate architectural features known as “network motifs”, i.e., the recurring patterns of gene interactions within the regulatory networks (Alon, 2021). Each network motif performs a defined information processing function within the network and some of them, such as the double negative feedback loop, can lead to “toggle switches” between different cell states (Alon, 2007). Thus, delineating the circuitry of interactions among different TFs is key to understanding the molecular mechanisms that govern cell fate decisions.

Among the key regulatory components, *Oct4*, *Sox2*, and *Nanog* constitute the core pluripotency TFs. They act by activating the expression of other pluripotency-associated factors and simultaneously repressing lineage-specific genes, while also upregulating their own gene expression and that of each other (Hackett and Azim Surani, 2014; Yeo and Ng, 2013). While *Oct4* and *Sox2* are fundamental for the formation of the inner cell mass (ICM) and the maintenance of the pluripotent state of mESCs, *Nanog* is important for the acquisition of pluripotency but dispensable once it is achieved (Chambers et al., 2007). Interestingly, although *Nanog* is expressed in mESCs and in the late post-implantation like epiblast stem cells (EpiSCs), its expression is transiently downregulated as cells differentiate to EpiLCs (Gu et al., 2016; Hackett and Azim Surani, 2014; Hayashi et al., 2011). Conversely, other TFs are necessary for a proper induction of formative pluripotency. Although both *Oct4* and *Otx2* are expressed in naïve mESCs, their reorganization to new enhancers has been shown to be required for the correct acquisition of a formative identity (Buecker et al., 2014). In addition, *Otx2* as well as *Foxd3* promote the transition to formative pluripotency by coordinating the silencing of naïve genes and the activation of early post-implantation epiblast markers (Acampora et al., 2017; Bleckwehl et al., 2021). Interestingly, as is the case of *Oct4, Otx2*, and *Foxd3*, most of the TFs shown to be important in the formative GRN and are also expressed in mESCs.

Here, we show that the TF *Oct6* is not expressed in ground-state mESCs but is rapidly induced as cells transition to formative pluripotency. *Oct6* is a well-established marker of formative and primed pluripotency (Buecker et al., 2014) and is important during the neural differentiation of mESCs (Zhu et al., 2014). However, its role in the transition to EpiLCs has not been extensively addressed. We show that the absence of Oct6 prevented the correct dismantling of the naïve TF circuitry in differentiating cells, partially blocking the morphological transformation observed in EpiLCs. Furthermore, premature expression of *Oct6* in naïve cells was sufficient to induce a morphology akin EpiLCs, while inducing formative genes like *Sox3*, *Dnmt3A*, *FGF5* and repressing *Nanog* in a bistable manner. Our results suggest that *Oct6* promotes the exit from naïve pluripotency through the repression of *Nanog* and the induction of formative specific genes. Collectively, our results show that *Oct6* and *Nanog* form a double negative feedback loop which could act as a toggle switch important for the transition to the formative state.

## Results

### 1. Oct6 is one of the earliest TFs induced during the exit from naïve pluripotency

To shed light on the conversion of GRNs from naïve to formative pluripotency, we initially focused on the TFs induced at early stages of differentiation, since they could be crucial to GRN restructuring. Based on a thorough study performed by Yang and collaborators (Yang et al., 2019), we identified TFs that were upregulated in the first 6 hours of the transition and remained highly expressed at 48 hours (Figure S1A). This preliminary analysis revealed known facilitators of formative pluripotency, such as *Otx2*, *Foxd3* and *c-Myc* (Acampora et al., 2017; Bleckwehl et al., 2021; Kim et al., 2020). Other TFs that exhibited this trend include the retinoic acid receptors *Rarg* and *Rxrg*, as well as *Myrf*, *Ar*, and *Aff3*. Interestingly, among these early-induced genes, *Oct6* (also known as *Pou3f1*) was the one that displayed the most important induction when comparing mESCs vs. 48h EpiLCs, and it was significantly upregulated only 2 hours after the onset of differentiation (Figure 1A). In contrast, other formative markers such as *Fgf5*, *Otx2*, and *Dnmt3A* were upregulated at later time points. The rapid induction of *Oct6* transcription was accompanied by a significant decrease of the naïve TF *Klf4* expression at 2h, whereas other naïve TFs such as *Nanog*, *Esrrb*, and *Tbx3* were downregulated at later time points. Quantitative immunofluorescence showed that OCT6 was not detected in naïve pluripotent ground state conditions and that its levels increased gradually as cells entered formative pluripotency, with all cells expressing this TF at 48 h of differentiation (Figure 1B, Figure S1B). In light of these results, we decided to assess the role of *Oct6* in the dissolution of naïve pluripotency.

**Figure 1.**
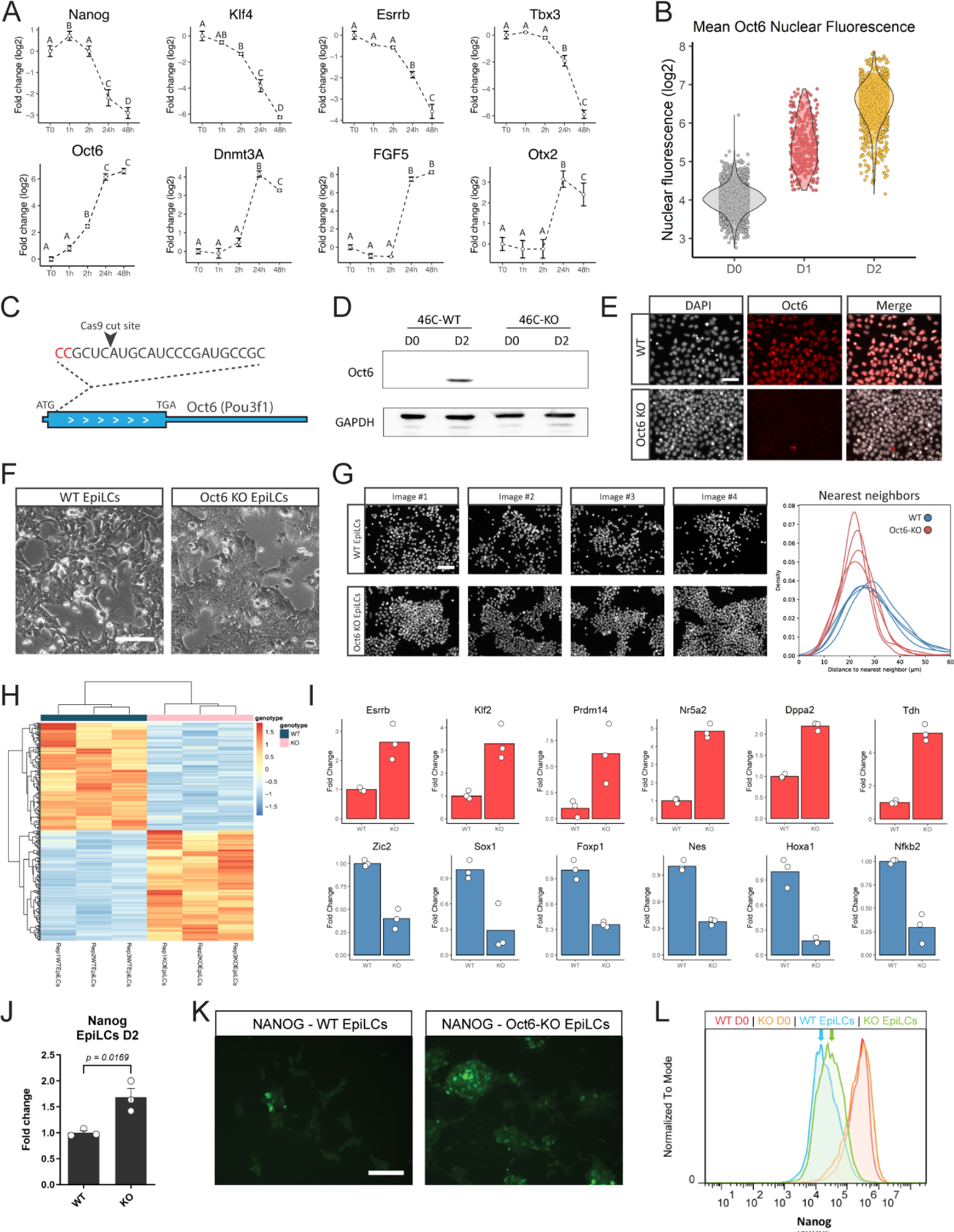
Loss of Oct6 affects the transition to formative pluripotency. (A) RT-qPCR analysis of naïve and formative pluripotency markers during the first hours of EpiLC differentiation. Results are presented as mean ± SEM for three independent replicates. Different letters indicate significant differences between time points (p<0.05). (B) Quantification of nuclear OCT6 during EpiLC induction. The violin plot shows the distribution of OCT6 nuclear intensity for each timepoint and the circles show the mean fluorescence for individual cells. (C) Diagram of Oct6 genomic locus showing the sgRNA binding region for CRISPR/Cas9 mediated KO generation. (D) Representative immunoblot showing OCT6 depletion in Oct6 KO EpiLCs. GAPDH was used as a loading control. (E) Representative immunofluorescence of OCT6 in WT and KO 48 h EpiLCs. DAPI was used as a nuclear stain. Scale bar: 50 µm. (F) Morphological differences between WT and Oct6 KO cells induced to differentiate to EpiLCs. Brightfield micrographs of WT and Oct6 KO EpiLCs. Scale bar: 50 µm. (G) *Left.* Representative images of DAPI stained nuclei of WT and Oct6 KO EpiLCs. *Right*. Distribution of the distances to the nearest neighbor cells for all nuclei in the representative images (see Materials and Methods). (H) Heatmap of differentially expressed genes between WT and Oct6 KO EpiLCs as obtained from Deseq2 analysis. (I) Examples of DE gene expression for naïve pluripotency genes (top) and formative genes (bottom) from the RNA-seq results. (J) RT-qPCR analysis of Nanog in WT and Oct6 KO EpiLCs. Results are as presented as mean ± SEM for three independent replicates. (K) Representative immunofluorescence of NANOG in WT and Oct6 KO EpiLCs. Scale bar: 100 µm. (L) Flow cytometry analysis of Nanog expression showing higher expression in Oct6 KO cells compared to WT EpiLCs.

We then conducted an analysis of the *Oct6* promoter by examining previously published ChIP-seq experiments to identify potential cis-regulatory elements (CRE) and likely regulators influencing Oct6 expression (Atlasi et al., 2019; Bleckwehl et al., 2021; Buecker et al., 2014; Chen et al., 2018; Narita et al., 2021; Yang et al., 2019). We discovered two CREs located 10 and 12 kb upstream of the transcription start site, which we refer to as CRE#1 and CRE#2, respectively (Figure S2). In mESCs maintained in 2i+LIF, the TFs OCT4, OTX2, NANOG, and ESRRB bind only to CRE#2 and not to CRE#1. Interestingly, EpiLCs presented a reorganization of OCT4 and OTX2 binding, with their occupancy extended to CRE#1. Moreover, this regulatory element exhibited an increased signal of the active enhancer marks H3K27ac and H3K4me1, along with open chromatin detected by ATAC-seq. These analyses collectively suggest that CRE#2 acts as a regulatory sequence specific to naïve pluripotency, potentially involved in repressing *Oct6* expression, while CRE#1 may function as an enhancer specific to EpiLCs.

### 2. Absence of Oct6 negatively affects the transition to formative pluripotency

To directly assess the role of *Oct6* in the transition to EpiLCs, we generated an *Oct6* knockout line (*Oct6-KO*) using CRISPR/Cas9 in 46C mESCs (Figure 1C, Figure S3A). Western blot and immunofluorescence analysis confirmed the lack of expression of OCT6 protein after induction to EpiLCs (Figure 1D,E). Importantly, KO cells cultured in ground state conditions did not show any changes in morphology or the expression of the pluripotency transcription factor *Nanog* (Figure S3B,C).

*Oct6* has been previously reported as a key player in neural progenitor cell (NPC) differentiation. We thus evaluated the ability of the *Oct6-KO* cells to differentiate to the neural lineage by taking advantage of 46C mESCs line, that expresses GFP under the control of the neural marker Sox1. As expected, *Oct6-KO* cells produced significantly lower rates of SOX1-GFP+ cells than WT 46C cells after 6 days of differentiation (∼60% vs. ∼90% SOX1-GFP+ cells, respectively) (Figure S3D). These data are consistent with *Oct6* promoting neural induction, although its expression is not fundamental for the differentiation of NPCs.

We then addressed if *Oct6-KO* cells were affected in their differentiation capacity to EpiLCs. The transition from naïve to formative pluripotency is accompanied by important morphological changes. mESCs in the naïve ground state grow as tightly packed colonies with a dome shape. Upon EpiLC induction, cells quickly undergo a morphological conversion that includes flattening, diminished cell-cell interactions, and the formation of cellular protrusions (Buecker et al., 2014; Waisman et al., 2019). Interestingly, we noticed that upon differentiation, *Oct6-KO* cells exhibited a more compact morphology with considerably fewer protrusions compared to parental WT cells (Figure 1F). Indeed, quantification of the distribution of distances to each cell’s nearest neighbor both for WT and KO EpiLCs confirmed that *Oct6-KO* cells were more tightly packed and did not show colony-detached cells as in the case of WT EpiLCs (Figure 1G). Thus, the absence of *Oct6* impairs the phenotypic changes observed as mESCs exit naïve pluripotency.

To further assess the role of Oct6 during differentiation we performed an RNA sequencing (RNA-seq) experiment on WT and *Oct6-KO* EpiLCs. A total of 292 genes were differentially expressed (DE) with at least 1 log 2-fold change (Figure 1H, Table S1). Functional annotation analysis of the DE genes with Gene Ontology (GO) revealed significant enrichment for the biological process term “regulation of cell motility” (GO:2000145) and the cellular component terms “plasma membrane bounded cell projection” (GO:0120025) and “cell leading edge” (GO:0031252), in agreement with the phenotypic effects previously observed. Importantly, the GO term “cell differentiation” (GO:0030154) was also highly enriched, suggesting that KO EpiLCs may display alterations in their differentiation capacity. Among the DE genes, we found that key TFs associated with naïve pluripotency such as *Esrrb, Klf2, Nr5a2, Dppa2, Tdh, Zic3*, and *Prdm14* showed higher expression levels in KO cells compared to WT EpiLCs (Figure 1I, Figure S3E). Interestingly, although not detected in the RNA-seq data due to the stringency of the analysis, assessment of the master pluripotency regulator *Nanog* also showed a slight but significant upregulation in KO EpiLCs, both at the mRNA and protein level (Figure 1J-L). On the other hand, genes associated with the transition to formative pluripotency or neural differentiation such as *Zic2*, *Sox1*, *Nestin*, *Foxp1*, and *Hoxa1* (Ying et al., 2003) were less expressed in KO EpiLCs. Supporting the role of Oct6 in the changes in cell morphology, we also observed differences in genes associated with cell attachment such as *Vimentin*, *E-Cadherin*, *N-Cadherin* and *Claudins 5, 6, 7* and *9* (Figure S3E,F). In summary, our results indicate that the absence of Oct6 impairs the correct acquisition of the GRN associated with formative pluripotency.

We next intended to assess the genome-wide binding of OCT6 in EpiLCs. However, there are currently no suitable *Oct6* antibodies for ChIP (see the Discussion). Therefore, we analyzed previously published data, where Matsuda et al evaluated OCT6 binding in epiblast stem cells (EpiSCs) by overexpressing a tagged version of *Oct6* (Matsuda et al., 2017). Although EpiLCs and EpiSCs represent different developmental stages of post-implantation epiblast development, we reasoned that these data could be useful to infer which of the DE genes in our RNA-seq experiment could be direct targets of OCT6. Indeed, among the 292 DE genes detected in our RNA-seq, 114 were associated with OCT6 ChIP binding peaks in EpiSCs (Figure S4A). This number was almost three times higher than the ∼40 genes that would be expected out of chance, as evaluated by a bootstrap analysis. Interestingly, among the DE genes of our RNA-seq that contained OCT6 peaks in EpiSCs we found the naïve pluripotency expressed genes *Dppa2, Prdm14, Nr5a2*, and *Vim* and the EpiLCs induced genes *Zic2, Zic3, Sox1, Nes,* and *Foxp1* (Figure S4B). Overall, this analysis validated our RNA-seq results and identified genes that are potentially regulated by *Oct6* at the transcriptional level by direct binding to their loci.

### 3. Overexpression of *Oct6* in naïve cells represses *Nanog* and induces morphological changes associated with formative cells

To further study the role of *Oct6* in the transition from naïve to formative pluripotency, we analyzed the effect of its premature expression in the pluripotent ground state. To that end, we engineered a new cell line where KO cells were complemented with a construct that allows the doxycycline (Dox) inducible expression of an HA-tagged version of OCT6, together with the fluorescent protein mCherry via a self-cleaving peptide (Figure 2A). As expected, the addition of Dox induced the expression of the *Oct6* and *mCherry* transgenes, while allowing the *in vivo* observation of red fluorescence in OCT6 overexpressing cells (Figure 2B). Quantitative immunofluorescence confirmed that mCherry signal was a good proxy of OCT6 expression since their levels were highly and linearly correlated throughout several orders of magnitude (Figure 2C). Importantly, induction of transgenic OCT6 expression in differentiating KO cells rescued the morphological effect observed previously, as seen by the reappearance of diminished cell-cell interactions and cell protrusions (Figure 2D).

**Figure 2.**
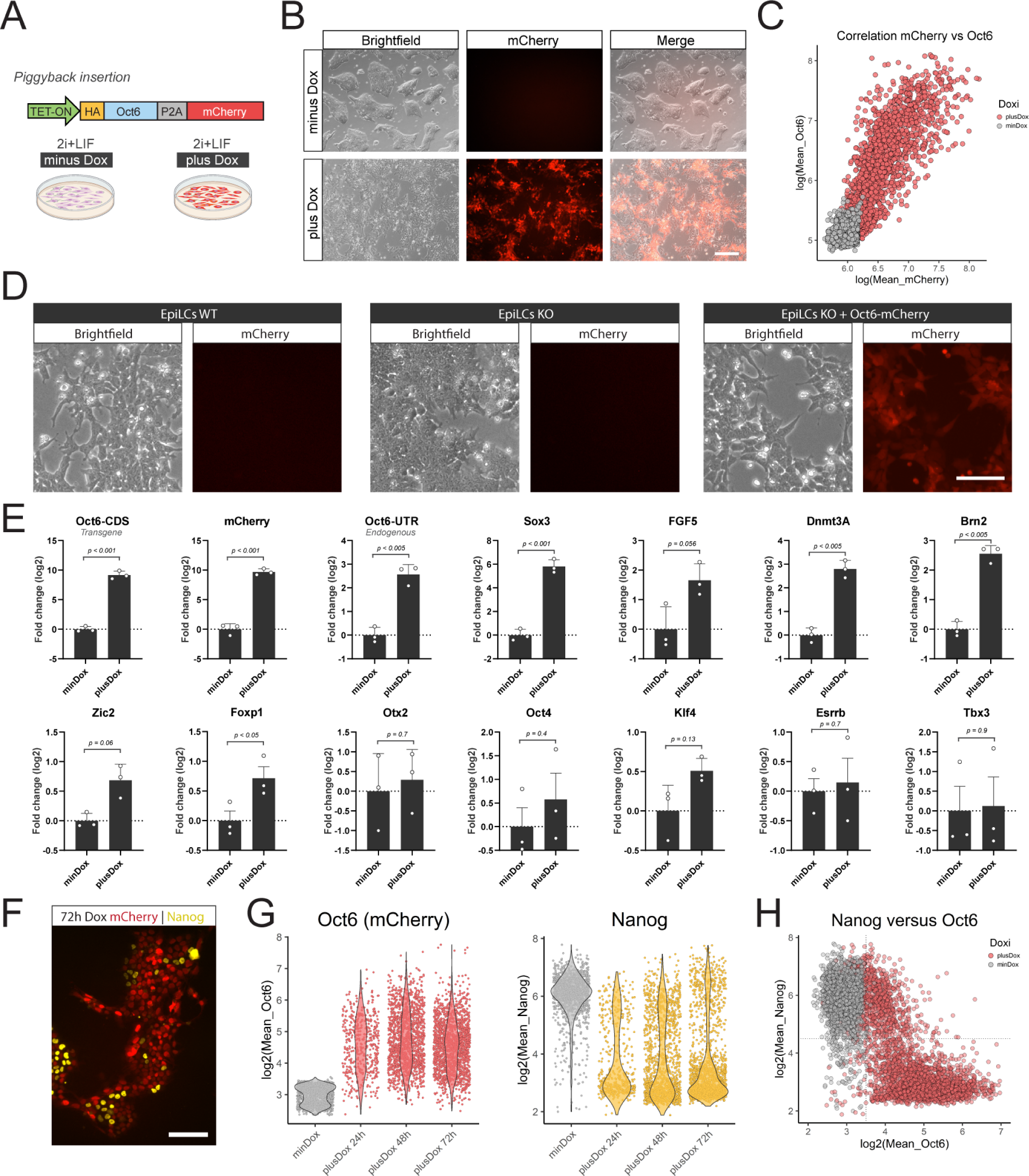
Overexpression of Oct6 in undifferentiated cells promotes morphological and transcriptional changes associated with formative pluripotency. (A) Diagram of the Oct6-P2A-mCherry overexpressing construct and the experimental design. (B) Treatment with Dox in 2i+LIF medium induces the expression of mCherry and a morphological transformation reminiscent of formative cells. Scale bar: 200 µm. (C) The plot shows the linear correlation of mean nuclear intensity fluorescence of mCherry and OCT6 obtained from OCT6 immunostaining of Dox treated cells. Each circle shows the data of one individual nucleus. (D) Overexpression of Oct6 in KO EpiLCs rescues the morphological differences observed between WT and KO EpiLCs. Scale bar: 50 µm. (E) RT-qPCR analysis of naïve and formative markers upon Dox treatment. Results are as presented as mean ± SEM for three independent replicates. (F) Representative immunofluorescence showing how Dox treatment induces the repression of NANOG in Oct6-P2A-mCherry expressing cells. Scale bar: 100 µm. (G) Quantification of immunofluorescence experiments of mCherry and NANOG in untreated or 24, 48, and 72 h Dox treated cells in 2i+LIF medium. The violin plots show the distribution of mCherry or NANOG nuclear intensity for each time point. The circles show the mean fluorescence for individual cells. (H) The plot shows the correlation between Oct6-P2A-mCherry and NANOG nuclear intensity expression in untreated or Dox-treated cells in 2i+LIF after immunostaining experiments. The circles show the mean fluorescence for individual cells. Data for 24, 48, and 72 h Dox treatment was pooled since they behaved similarly.

We next evaluated the effect of *Oct6* expression in ground state conditions. While dox untreated cells formed typical compact dome-shaped colonies, the addition of Dox for 72 h induced a pronounced morphological change similar to the observed upon differentiation, with flattened colonies, diminished cell-cell interactions, and the formation of cellular protrusions (see Figure 2B). These changes could be observed as early as 24 h after Dox treatment. As an additional control, no morphological changes were observed when using an inducible cell line that only overexpressed *mCherry* (Figure S5A). Gene expression analysis showed that induction of *Oct6* expression in 2i+LIF media upregulated the formative marker genes *Sox3*, *Dnmt3A, Fgf5, Foxp1, Brn2, Zic2, Zic3*, and endogenous *Oct6*, while not affecting *Otx2* nor the general pluripotency marker *Oct4* (Figure 2E, Figure S5B). Of the formative marker genes, only *Dnmt3A* lacks OCT6-binding peaks in EpiSCs, suggesting that the remaining genes could be direct targets of *Oct6* (see Figure S4B). Importantly, no significant changes were observed in the expression of the naïve pluripotency markers *Esrrb, Rex1, Prdm14 Klf4*, and *Tbx3*.

To gain a deeper insight into the regulatory effect of Oct6, we performed quantitative immunofluorescence experiments against key TFs regulated in this transition. OCT4, SOX2, and KLF4 showed similar protein levels between Dox treated and untreated mESCs, with only a slight reduction in the mean expression (Figure S5C). In the case of SOX3, consistent with the gene expression data, we detected an important upregulation in a subset of OCT6 overexpressing cells, further reinforcing the regulatory link between these genes. We were particularly interested in the analysis of *Nanog*, not only because of its increased transcript levels in KO versus WT EpiLCs, but also because we discovered that NANOG and OCT6 proteins are expressed in a mutually exclusive manner in WT EpiLCs (Figure S6A). Strikingly, while more than 90% of untreated cells expressed high levels of NANOG, we observed that overexpression of OCT6 repressed the expression of NANOG at the protein level in more than 70% of the cells, both at 24, 48 and 72 h of Dox (Figure 2F, Figure S6B). This effect was positively correlated with OCT6 levels and displayed a bistable switch-like behavior (Figure 2G). As a result, cells that were mCherry negative were more than 80% NANOG positive, while mCherry expression two-fold above background levels already showed more than 40% NANOG negative cells, suggesting that mild OCT6 expression is sufficient to repress NANOG (Figure 2H). For cells expressing mCherry at levels beyond four-fold above background, more than 90% of cells were NANOG negative. Interestingly, the fact that NANOG exhibited a clear bistabe “ON-OFF” expression might indicate the existence of a repressive feedback loop between these TFs.

We next wondered whether the repression of NANOG was reversible after releasing the induction of OCT6. To answer this, we removed Dox from 72 h treated cultures and analyzed mCherry and NANOG protein levels 24, 48, and 72 h after washing the cells (Figure S7A). Interestingly, as early as 24 h after Dox removal, mCherry fluorescence returned to background levels and NANOG expression was restored (Figure S7B). We did not observe dead cells in these cultures, indicating that NANOG+/OCT6-derived from previously NANOG-/OCT6+ cells. The morphological changes induced by Dox were also reverted and colonies re-acquired their typical highly packed dome shape. Overall, these results indicate that OCT6 expression in naïve culture conditions induces a reversible non-physiological identity with similarities to the formative pluripotent state.

### 4. OCT6 represses *Nanog* at the transcriptional level

To further analyze the dose dependency of OCT6 in NANOG repression, we took advantage of the expression of mCherry and analyzed 24 h Dox-treated cells by flow cytometry, gating them into 3 populations, Low, Med, and High, all of them with the same number of cells (Figure 3A). In agreement with our previous results, ∼75% of cells in the mCherry-low population displayed high NANOG levels, while these percentages decreased to ∼35% and ∼2% in the mCherry-Med and mCherry-High populations, respectively (Figure 3B). By sorting these 3 cell populations, we confirmed the correlation between mCherry fluorescence and the mRNA levels of *Oct6*, *mCherry*, and the endogenous *Sox3* mRNAs (Figure 3C). Interestingly, mCherry expression showed an inverse relation with *Nanog* mRNA levels, suggesting that OCT6 repressed *Nanog* at the transcriptional level. Even though the mCherry-Med population contained 65% of cells with reduced NANOG protein levels, we only observed a roughly two-fold reduction in its mRNA levels. This could be explained by the contribution of the remaining ∼35% fully NANOG positive cells, which can mask the downregulation when analyzing the mean Nanog mRNAs levels in the entire population. To complement these findings, we conducted a luciferase reporter assay to assess the impact of OCT6 expression on the Nanog promoter. Remarkably, induction of Oct6 using Dox resulted in a significant decrease in Nanog reporter activity, observed in both undifferentiated and 24 h differentiating cells (Figure S8A). These findings provide compelling evidence of OCT6’s regulatory influence on the Nanog promoter, further supporting the notion of regulation at the transcriptional level. To reinforce these results, we performed single-molecule RNA-FISH experiments to detect individual *Nanog* mRNAs in OCT6 overexpressing cells. We first validated this method by analyzing undifferentiated mESCs and 24 h EpiLCs, which confirmed wide expression of *Nanog* transcripts in naïve cells and an important reduction as cells transited towards the formative state (Figure S8B). After this validation, we evaluated the distribution of *Nanog* transcripts in undifferentiated cells induced to express *Oct6* while simultaneously detecting NANOG and OCT6 proteins. As expected, cells with low levels of OCT6 protein were positive for *Nanog* transcripts and expressed NANOG at the protein level. Interestingly, cells with high levels of OCT6 protein did not present neither NANOG protein nor *Nanog* mRNAs (Figure 3D). In summary, our results demonstrate that OCT6 expression in naïve ground state conditions represses *Nanog* at the transcriptional level. Together with the over-expression of formative-specific transcription factors such as *Sox3* and the *de novo* DNA methyltransferase *Dnmt3A*, *Oct6* might ultimately induce a shift in the naïve GRN that partially sets it to a formative configuration similar to the early post-implantation epiblast.

**Figure 3.**
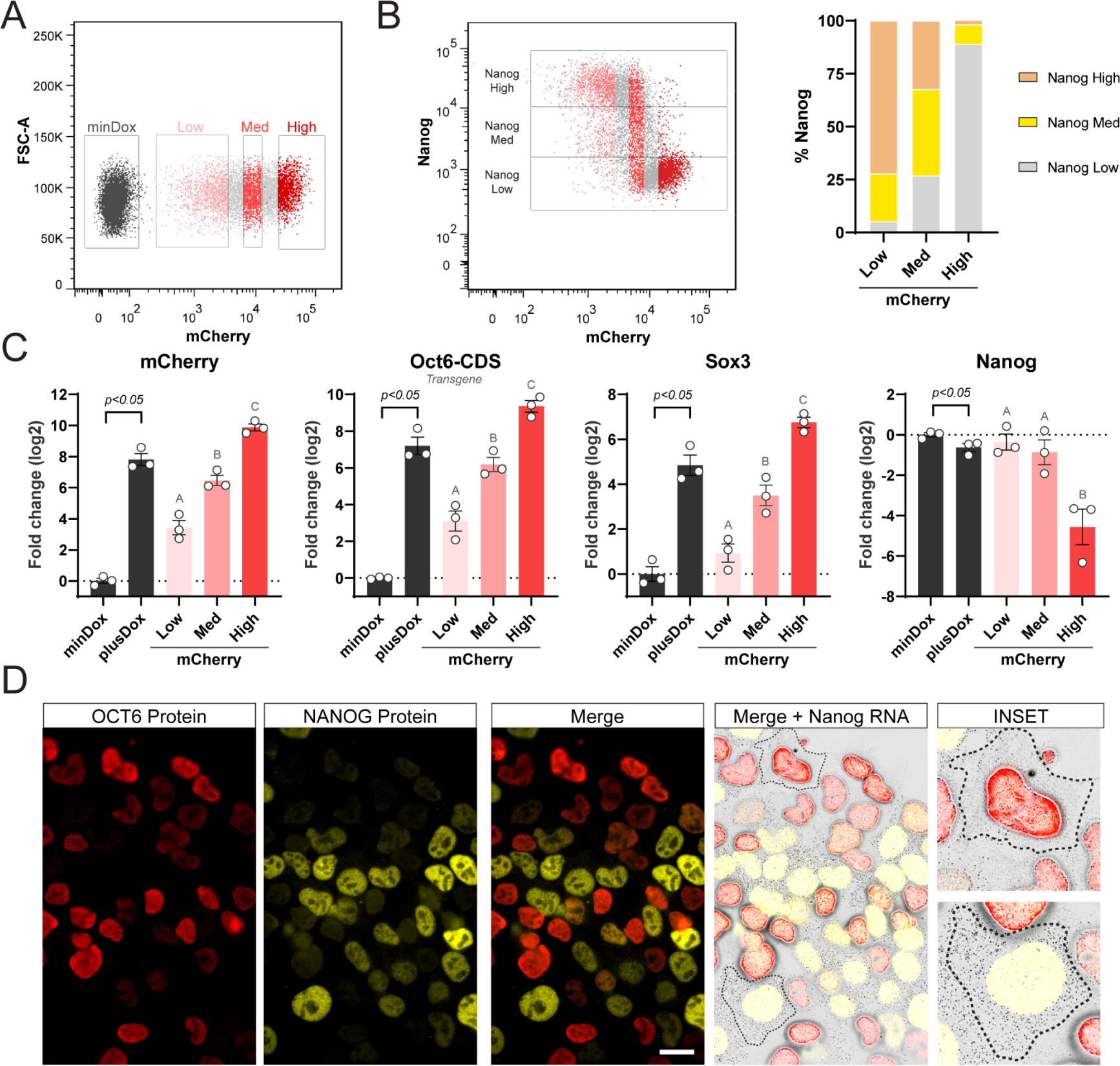
Oct6 represses Nanog at the transcriptional level. (A) Flow cytometry sorting of 24 h Dox treated cells into populations with low, medium (med), and high levels of mCherry. (B) *Left*. Correlation between mCherry and NANOG by flow cytometry. The different mCherry expressing subpopulations are shown. *Right*. The proportion of NANOG high, med, and low in the different mCherry expressing subpopulations. (C) RT-qPCR analysis of exogenous *Oct6*, *mCherry*, *Sox3*, and *Nanog* in *Oct6* overexpressing cells. Untreated or Dox-treated cells (bulk population) are shown in black and statistically compared using a student’s t-test. The different mCherry expressing subpopulations are shown in shades of red and were compared using random block ANOVA. Different letters indicate significant differences (p<0.05). (D) Simultaneous single-molecule RNA FISH against Nanog cytoplasmic transcripts and immunostaining against Nanog and Oct6 proteins. The insets show example cells with high and low Oct6 transgene expression. Scale bar: 20 µm.

### 5. *Oct6* and *Nanog* form a double negative feedback loop with positive autoregulation

Our results so far demonstrated that OCT6 and NANOG are expressed in a mutually exclusive manner in WT EpiLCs and that OCT6 activates its own transcription while repressing the expression of Nanog. Moreover, Nanog is known to present positive autoregulation (Boyer et al., 2005; Saunders et al., 2013). This led us to hypothesize that these two TFs could constitute a double negative feedback loop that could act as a toggle switch to initiate the dissolution of the naïve pluripotent state. Indeed, we have previously shown that NANOG binds to CRE#2 in *Oct6’s* promoter along with OCT4, OTX2 and ESRRB (see Figure S2). To assess if OCT6 might also bind to the *Nanog* locus, we analyzed the aforementioned work by Matsuda et al in EpiSCs. While the ChIP-seq signal in *Nanog* locus exhibited relatively low intensity, we observed a discrete accumulation of reads at the 5 kb distal enhancer located upstream of the Nanog TSS (Figure S8C). This distal enhancer, recognized as a crucial CRE, is known to be targeted by other key pluripotency transcription factors (Blinka et al., 2016; Levasseur et al., 2008). This subtle observation prompted us to consider that OCT6 might bind to Nanog’s promoter in EpiLCs.

Finally, we evaluated if NANOG inhibited the expression of *Oct6* by generating a cell line capable of expressing a *Nanog* transgene under the control of Dox (Figure 4A). As in the case of *Oct6* overexpression, this cell line also expresses mCherry via a self-cleaving peptide. To confirm the correct behavior of this line, we first differentiated them for 48 h in the presence or absence of Dox and confirmed that they expressed mCherry and NANOG in a highly correlated fashion (Figure 4B,C). Next, we evaluated whether Dox-treated EpiLCs showed reduced levels of OCT6. Our results show that while mCherry negative cells exhibited normal levels of OCT6, mCherry expressing cells did not express this transcription factor, indicating that NANOG repressed *Oct6* in a cell-autonomous fashion (Figure 4D,E). Overall, our results indicate that these genes could constitute a toggle-switch circuit important for the correct dissolution of naïve pluripotency and the transition to a post-implantation epiblast-like phenotype.

**Figure 4.**
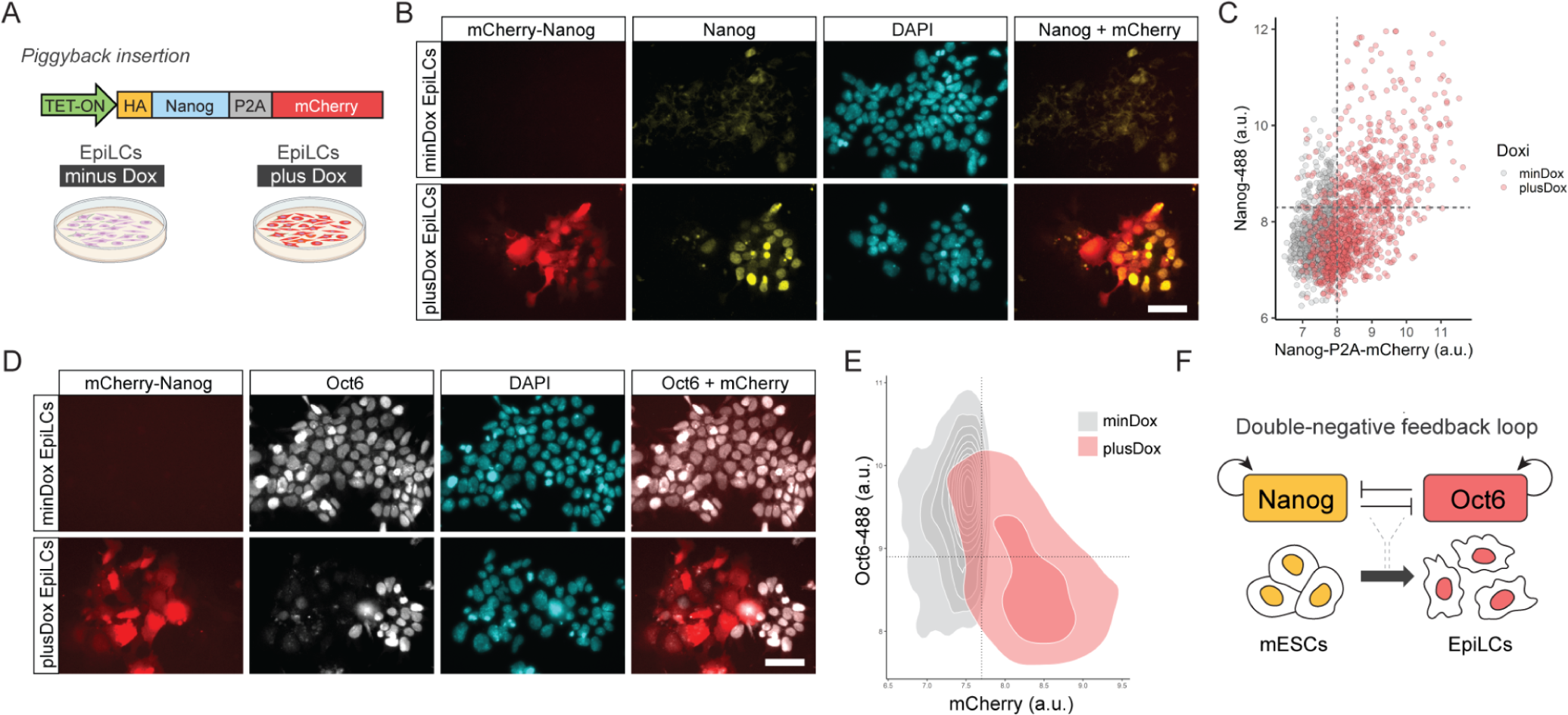
Oct6 and Nanog repress each other forming a double negative feedforward loop network motive. (A) Diagram of the Nanog-P2A-mCherry overexpressing construct and the experimental design. (B) Treatment with Dox during EpiLC differentiation induces the expression of mCherry. Scale bar: 50 µm. (C) The plot shows the correlation of mean nuclear intensity fluorescence of mCherry and NANOG obtained from NAONG immunostaining of Dox-treated EpiLCs. Each circle shows the data of one individual nucleus. (D) Immunostaining of OCT6 showing that Dox-treated Nanog-P2A-mCherry overexpressing cells do not express OCT6. Conversely, Dox-treated cells not expressing Nanog-P2A-mCherry normally express OCT6. Scale bar: 50 µm. (E) The plot shows the correlation of mean nuclear intensity fluorescence of Nanog-P2A-mCherry and OCT6, obtained from OCT6 immunostaining of Dox-treated EpiLCs. Each circle shows the data of one individual nucleus. (F) Proposed model of transcriptional circuitry between Nanog and Oct6 that regulates the transition from naïve to formative pluripotency.

## Discussion

In this study, we identified OCT6 as an important TF in the regulation of the formative pluripotent state as well as in dismantling the previous naïve identity. OCT6 belongs to the POU domain family of TFs, which are known to play critical roles in embryonic development and cellular differentiation. Previous studies have implicated *Oct6* in various biological processes, including neural development, axon regeneration, and glial cell differentiation (Kuhlbrodt et al., 1998; Mistri et al., 2015). Recently, it was shown that *Oct6* can replace *Oct4* in reprogramming human cells but fails to do so in mouse cells (Kim et al., 2020). Whether this difference relies on the different types of pluripotency presented by human and mouse PSCs, i.e., naïve vs. primed, remains to be fully investigated. Indeed, *Oct6* has recently been shown to block the differentiation of porcine IPSCs, which display a primed pluripotency phenotype (Yang et al., 2022). However, its precise role in the context of mouse PSC differentiation and the dissolution of naïve pluripotency circuitry remains largely unexplored.

Here, to identify key effectors of the conversion from naïve to formative pluripotency, we first searched for TFs that were rapidly induced and displayed high expression throughout the first 48 h of differentiation, when formative identity is established. Using this strategy, we identified Oct6 as a gene that is induced as early as 2 h after the onset of differentiation, with its expression peaking at 24 and 48 h of EpiLC induction. We show that OCT6 protein is expressed homogeneously in EpiLCs, reaching its maximum at 48 h. Importantly, OCT6 was not detected in ground state pluripotency when cells are cultured under defined conditions (N2B27+2i+LIF). This finding challenges previous reports that indicate that OCT6 can be detected in naïve cells (Suzuki et al., 1990; Zhu et al., 2014). This inconsistency might be attributed to the culture method, as mESCs cultured in undefined conditions containing serum result in heterogeneous populations where naïve cells coexist with primed and differentiated cells (Gu et al., 2016; Kolodziejczyk et al., 2015). OCT6 expression in these culture conditions could thus be restricted to the primed pluripotency compartment. Additionally, we discovered that several commercial antibodies are not specific for OCT6 and may detect other members of the Pou family of TFs (data not shown), underscoring the need for careful assessment of OCT6 expression. Overall, the fact that *Oct6* is induced during the transition to the formative state is consistent with its expression being first detected at the egg-cylinder stage of the mouse embryo (Zwart et al., 1996).

We then investigated the function of *Oct6* during the transition to formative pluripotency by generating an Oct6 KO mESC line. The absence of OCT6 resulted in cells organized as densely packed colonies more reminiscent of the naïve phenotype, as opposed to the typical formative morphology characterized by reduced cell-cell interactions and protrusions (Buecker et al., 2014; Waisman et al., 2019). Indeed, KO cells showed altered expression of genes associated with motility and cell morphology, such as *Vim*, *E-Cad*, *N-Cad*, and *claudin* proteins. Importantly, transcriptome analysis revealed that key naïve TFs, including *Nanog, Klf2, Esrrb, Prdm14, Nr5a2, Dppa2,* and *Tdh* (Du et al., 2010; Hackett and Azim Surani, 2014; Kolodziejczyk et al., 2015), were not properly downregulated in KO cells, indicating inefficient dismantling of the naïve GRN circuitry. We also observed reduced expression of genes normally induced in EpiLCs, such as *Zic2* and *Foxp1* (Hackett et al., 2018; Smith and Smith, 2014), and early neural genes weakly expressed in EpiLCs, including *Nes* and *Sox1*. Collectively, our findings support a critical role for *Oct6* in the transition from naïve to formative pluripotency.

Other groups have recently analyzed the effect of reducing OCT6 levels during mESCs differentiation, either by knockout or knockdown strategies. Kalkan and collaborators observed that *Oct6* KO mESCs did not delay the exit from naïve pluripotency (Kalkan et al., 2019); however, this was only evaluated by the expression of a *Rex1* reporter at 26 h of differentiation. Consistent with this, we did not observe differences in the expression of *Rex1* when comparing WT and KO 48 h EpiLCs. On the other hand, Zhu and collaborators showed that *Oct6* is important for the neural differentiation of mESCs specifically during the transition from epiblast stem cells (EpiSCs) to NPCs (Zhu et al., 2014). The impairment in neural differentiation was later confirmed by another group using *Oct6* KO mESCs (Li et al., 2017). Our results are consistent with this work, as *Oct6* KO cells only generated 60% SOX1-GFP+ cells compared to 90% generated by the parental WT cell line. However, the fact that the formative state was affected in KO cells and that the naïve circuitry was not completely dismantled could also explain the reduced efficiency of neural differentiation.

An important limitation of our study was the inability to perform genome-wide binding analysis of OCT6 in EpiLCs, as there are no suitable antibodies for chromatin immunoprecipitation (ChIP). Despite our efforts to evaluate various antibodies we were unable to obtain specific signals when analyzing different genomic locations. We also attempted the Cut and Run technique (Kaya-Okur et al., 2019) with our engineered cell line expressing HA-Oct6, using both OCT6 and HA-specific antibodies, but unfortunately, specific signals were not detected. Interestingly, Matsuda et al. successfully captured OCT6 binding to the genome by overexpressing a tagged version of this TF in EpiSCs, a slightly more advanced stage of postimplantation epiblast development (Matsuda et al., 2017). By comparing their data with our RNA-seq analysis, we found that 39% of the differentially expressed genes between WT and KO EpiLCs presented OCT6 binding peaks, a proportion significantly higher than the 14% expected by chance. Although this analysis strongly validates our results, future research should address diverse strategies to map the binding of OCT6 to the genome in EpiLCs.

The absence of OCT6 had a significant impact on the transition to formative pluripotency, although it did not completely hinder the dissolution from the naïve state. Interestingly, Oct6 KO mice are not embryonic lethal and only die within the first few days after birth (Jaegle et al., 1996), suggesting the presence of compensatory mechanisms during early development. To investigate the specific role of Oct6 within the pluripotency GRN, we examined the effects of its premature expression in the naïve state. Our findings demonstrate that expression of Oct6 in 2i+LIF media induces a morphological change similar to that observed in WT EpiLCs, while activating formative genes such as Sox3, FGF5, Dnmt3A, and the endogenous Oct6. Notably, Sox3 and FGF5 contained OCT6 binding peaks in EpiSCs, indicating their potential direct regulation by this TF. Strikingly, the majority of cells overexpressing OCT6 repressed the expression of the core pluripotency gene Nanog, evidenced at both the transcriptional and protein levels, without affecting OCT4, SOX2, and KLF4. This finding is noteworthy, as downregulation of Nanog is necessary to allow the exit from the naïve ground state (Abranches et al., 2013; Chambers et al., 2007; Liu et al., 2016). Although OCT6 overexpression displayed high heterogeneity and a wide range of expression, we observed that OCT6 levels immediately above background were sufficient to repress *Nanog*. Furthermore, we observed a subtle ChIP-seq signal of OCT6 binding to Nanog’s distal enhancer in EpiSCs, indicating that this TF might directly bind Nanog regulatory regions in EpiLCs. Interestingly, NANOG exhibited a bistable “ON-OFF” behavior within the “low OCT6” population, which is characteristic of double negative feedback loop subcircuits. To study this possibility, we evaluated Nanog and Oct6 expression in WT EpiLCs, which showed that these two proteins were expressed in a mutually exclusive manner. This prompted us to study whether NANOG also repressed *Oct6*. Our analysis of published data revealed that NANOG binds to the *Oct6* promoter at a CRE regulated by other key TFs that potentially inhibits Oct6 expression in the undifferentiated state. Importantly, we observed that overexpression of NANOG in differentiating cells blocked the induction of OCT6, consistent with a recent study (Barral et al., 2019). By causally demonstrating the mutual repression between Nanog and Oct6, along with their positive autoregulation, we propose that these two TFs form a double negative feedback loop (Figure 4F), serving as a toggle switch crucial for the transition to the formative state and providing robustness to the GRN of naïve and formative pluripotency.

## Authors Contributions

AW conceived, performed, and analyzed most of the experiments. DS and AW constructed the Nanog overexpressing cell line and analyzed the overexpression of Nanog in differentiating cells. AW and FS performed the Oct6 Dox reversion experiments and helped with many immunofluorescence experiments. MF performed the luciferase reporter experiments. AL, CB, BP, and GA contributed to proteomic and immunofluorescence analyses. ALG, AS, JS, and AW oversaw bioinformatic analyses. LM, GS, and AG were fundamental in their contribution to useful discussions. AW and SM discussed all the experiments and wrote the manuscript.

## Acknowledgements

We would like to thank Dr. Hernán Grecco and his lab for their help with single-molecule RNA-FISH experiments. We are also grateful to Alejandra Ventura and Gabriel Neiman for the important discussions that contributed to this work. The following grants provided financial support to this work: FONCYT: PICT-2011-1927, PICT-2015-1469, and PID-2014-0052; CONICET: PIP2015-2017, PICT-2018-02672, PICT-2018-00836, PICT-2018-01722, PICT-2020-03553, PICT2020-03523. We would also like to thank Fundación FLENI and Fundación Perez Companc for their continuous support.

## Star Methods

### Cell Culture

46C mESCs and the transgenic lines derived from them were cultured using the chemically defined medium N2B27 with 1000 U/ml LIF (Millipore), 1 µM PD0325901 (Tocris) and 3 µM CHIR99021 (Tocris), hereafter called ‘2i+LIF medium’. N2B27 formulation is a 1:1 mix of DMEM-F12 (Invitrogen Cat# 12634) and neurobasal medium (Invitrogen Cat# 21103), supplemented with N2 (Invitrogen Cat# 17502), B27 supplement without Vitamin A (Invitrogen Cat# 12587), 2 mM GlutaMAX (Invitrogen Cat# 35050), 40 μg/ml of BSA (Invitrogen Cat# 15260), 10 μg/ml insulin (Sigma-Aldrich Cat# I9278), 0.1 mM β mercaptoethanol (Invitrogen Cat# 21985). Cells were maintained on 0,1% gelatin-coated dishes, passaged every 2-3 days using TrypLE (Gibco) and cultured at 37°C in a 5% CO2 (v/v) incubator. Cells were cultured without antibiotics and routinely assessed for mycoplasma contamination by PCR. To induce differentiation from the naïve ground state to EpiLCs, 50.000 cells/cm^2^ were plated in 15 µg/ml Fibronectin (Thermo Fisher Scientific) coated dishes in the “EpiLC” medium. This medium consisted of base N2B27 supplemented with 1% KSR, 12 µg/ml bFGF, and 20 µg/ml Activin A (all from Thermo Fisher Scientific). Cells were differentiated for 24 or 48h, as indicated in the main text. 46C mESCs were a kind gift from Dr. Austin Smith.

### Generation of Oct6 KO 46C mESCs

46C mESCs Oct6 knockout cell line was constructed using CRISPR/Cas9. We designed two specific sgRNAs targeting the 5’ region within the Oct6 coding sequence to generate an indel next to the initiation codon. sgRNAs were designed using CHOPCHOP V3 (Labun et al., 2019). Sense and antisense oligonucleotides were annealed and cloned into a BbsI-linearized pX330-GFP-Puro plasmid (Waisman et al., 2017). Recombinant plasmids were validated by DNA sequencing. Wild type 46C mESCs were independently transfected with the sgRNA plasmids using Linear Polyethylenimine (PEI, Polysciences) with a DNA/PEI ratio of 1:3 and selected with 3 μg/ml puromycin from 24 to 72 h post-transfection. After limiting dilution, 12 individual clonal lines were obtained for each sgRNA, amplified, and cryopreserved. Knockout generation for each clone was evaluated by genomic DNA sequencing of a PCR-amplified Oct6 locus. A clone with a homozygous insertion of a nucleotide within the Cas9 cutting site was chosen and further validated by Western Blotting and immunofluorescence, as indicated in the main text. Sequences for sgRNA sequences and KO evaluation primers are listed in Table S2.

### Generation of Oct6-P2A-mCherry, Nanog-P2A-mCherry and mCherry overexpressing cell lines

To overexpress Oct6 in an inducible manner we designed a piggyBac plasmid that could be integrated into the cell’s genome through a transposable recombination approach. The sequence coding for Oct6-P2A-mCherry was designed *in silico*, fully synthesized by SynBio Technologies, and cloned into a piggyBac target vector. This plasmid expresses the TET repressor together with a delta-puro antibiotic resistance gene. It also contains a TET-ON promoter followed by an AgeI and NotI restriction site where the Oct6 cassette was cloned. The HA-Oct6-P2A-mCherry cassette contains the following elements in the 5’-3’ orientation: an AgeI site followed by the Kozak sequence, an hemagglutinin (HA) tag fused to the Oct6 coding sequence in the amino termini, a HindIII site, the P2A self-cleaving peptide sequence, a MluI sequence and the mCherry coding sequence followed by a stop codon. This plasmid, named ePB-HA-Oct6-P2A-mCherry, was co-transfected into 46C Oct6 KO mESCs together with the piggyBac transposase at a ratio of 2:1, respectively, using PEI, as described previously. After puromycin selection for 10 days, clonal cell lines were generated by limiting dilution, amplified, and cryopreserved. To evaluate the expression of Oct6-P2A-mCherry cells were treated with Dox at concentrations ranging from 50 ng/ml to 2000 ng/ml. The concentration of 200 ng/ml of Dox and cell clone #3 was chosen as they produced the most homogeneous mCherry expression.

To generate the Nanog-P2A-mCherry cell line we followed a similar strategy. Briefly, the Nanog coding sequence was synthesized by SynBio Technologies and cloned into the piggybac plasmid constructed previously, between the AgeI and HindIII sites. WT 46C cells were co-transfected with this plasmid and the piggyBac transposase as previously described. Cells were treated with puromycin for 10 days to select stable transfectants. mCherry and Nanog expression was verified by Dox treatment at 200 ng/ml.

To generate the control mCherry inducibly expressed cell line, we modified the ePB-HA-Oct6-P2A-mCherry plasmid to remove the Oct6 coding sequence flanked by the AgeI and MluI sites. We exchanged this sequence with a short *in vitro* annealed sequence containing the AgeI and MluI overhangs and an ATG sequence to initiate the translation of the mCherry transgene. This plasmid was co-transfected into WT 46C mESCs with the piggyBac transposase as previously described. Cells were treated with puromycin for 10 days to select stable transfectants. mCherry expression was verified by Dox treatment at 200 ng/ml.

### Quantitative Real-Time PCR

Total RNA was extracted with Trizol (Thermo Fisher Scientific) following the manufacturer’s instructions, treated with DNAse (Thermo Fisher Scientific), and reverse transcribed using MMLV reverse transcriptase (Promega). Quantitative PCR was performed in a StepOne Real-Time PCR system (Applied Biosystems). Primer efficiency and N0 values were determined by LinReg software (Ruijter et al., 2009). Gene expression was normalized to the geometrical mean of GAPDH and PGK1 housekeeping genes, data was then log-transformed and relativized to the average of the biological replicates for the 2i+LIF condition. Statistical significance for qPCR data was analyzed by either Student-T test or randomized block design ANOVA. In the latter case, comparisons between means were assessed using the Tukey test. Plots were generated either with R or with Graphpad 8.

### Quantitative immunofluorescence

For immunofluorescence experiments, cells were grown on 12 mm coverslips coated for 30 minutes with Geltrex (Thermo Fisher Scientific). Cells were fixed for 15 minutes with 4% paraformaldehyde, permeabilized with 0.1% Triton X-100 PBS (PBST), and blocked with 3% normal donkey serum in PBST. Primary antibodies (Table S3) were added in block solution, incubated at 4°C overnight, and then washed three times in PBST for 30 minutes. Secondary antibodies and DAPI were incubated in block solution at room temperature for 30 minutes. Samples were washed as before, rinsed with distilled water, and mounted and imaged at 20X on an EVOS fluorescence microscope with adequate parameters (Thermo Fisher Scientific).

For quantifications, we followed a pipeline that included background subtraction, automatic nuclear segmentation using Stardist (Schmidt et al., 2018), and nuclear fluorescence intensity measurements for different channels with custom Fiji/ImageJ Macros. Measurement tables were then analyzed using custom R scripts with the tidyverse library. For nearest-neighbor analysis, the central nuclear position of each cell was obtained using Fiji/ImageJ macros and data was analyzed in Python with the scikit-learn library.

### Single Molecule RNA-FISH

smRNA-FISH was performed using Stellaris RNA-FISH probes (Biosearch Technologies) that we designed specifically to recognize Nanog mRNAs (Table S4). For smRNA-FISH alone (no immunofluorescence), we followed the manufacturer’s protocol for adherent cells. Briefly, cells were fixed with PFA 4%, permeabilized with cold ethanol 70% for 24h, and then treated 2 times for 3 minutes with FISH Wash Buffer. The wash buffer consisted of 2X sodium citrate buffer (SSC, Roche) and 5% deionized formamide (Sigma-Aldrich) in nuclease-free water. Nanog Stellaris FISH Probes Quasar® 670 were added to the hybridization buffer (HB) and incubated with coverslips containing fixed cells for 16 hours at 37°C. HB consisted of 0.1% dextran sulfate (Sigma-Aldrich), 2X SSC, and 10% deionized formamide in nuclease-free water. After incubations, cells were rinsed once with Wash Buffer and incubated with the same solution at 37°C for 30 minutes. After two rinses with nuclease-free PBS, nuclei were stained with DAPI and mounted into glass slides using Prolong Diamond Antifade Mountant (Thermo Fisher Scientific).

For immunofluorescence coupled with smRNA-FISH, we first followed the regular immunofluorescence protocol for the desired antigens. After washing the secondary antibodies, cells were re-fixed with 4% PFA for 15 minutes and the smRNA-FISH protocol was initiated and performed as described above.

FISH images were captured in an Axio Observer Z1 epifluorescence microscope (Zeiss) using an EC Plan-Neofluar 63x/1.25 Oil M27 and the ZEISS Axiocam 702 mono camera. Z-stacks encompassing the entire volume of the cells were taken with maximum Z resolution. For the smRNA-FISH channel, each Z-stack was exposed for 3 seconds with maximum LED intensity. After obtaining the images, a Laplacian of Gaussian transformation was applied to the smRNA FISH channel, and a maximum intensity projection (MIP) was generated and merged with the MIPs of the other fluorescence channels.

### Western blot analysis

Protein samples were collected with RIPA buffer, resolved on 10% SDS-polyacrylamide gels, and transferred to PVDF membranes (Amersham). Membranes were blocked for 1 h at room temperature in TBST 1% BSA and primary antibodies (Table S3) were incubated overnight at 4°C in block. Peroxidase-bound secondary antibodies were incubated for 1 h at room temperature. Membranes were revealed with ECL in a G-Box System (Syngene).

### Flow Cytometry

Flow cytometric analysis of unfixed Sox1+ neural precursor cells or fixed immunostaining against Nanog was performed in an Accuri C6 benchtop cytometer (BD). Immunostaining was performed with similar steps as for immunofluorescence, except that cells were previously trypsinized, fixed, and stained with primary and secondary antibodies in suspension for 1 h each. Cell sorting of mCherry+ live mESCs was performed on a BD FACS Aria FUSION flow cytometer. Sorted populations were immediately treated with Trizol reagent for posterior RNA extraction.

### Analysis of the Nanog-Luciferase reporter

To evaluate firefly luciferase activity in response to Oct6 expression, Nanog5P reporter (Addgene #16337) was co-transfected together with the ePB-HA-Oct6-P2A-mCherry plasmid into undifferentiated mESCs. Transfected cells were either maintained in this state or differentiated for 24 h, either with or without Dox. Luciferase activity was analyzed using as previously reported (Francia et al., 2021).

### RNA-seq and bioinformatic analysis

For the analysis of TFs induced during the early stages of EpiLC differentiation, processed RNA-seq data from Yang et (GSE117896) were downloaded from the GREIN database (Mahi et al., 2019). Raw data was fed to the into DESeq2 for differential expression analysis (fdr<0.1) between 0 h and 6 h of EpiLC induction. Among the upregulated genes, TFs were selected using the Biomart package in R, filtering by the GO term GO:0003700 “DNA-binding transcription factor activity”.

For the RNA-seq experiments performed in this work, WT and Oct6 KO cells were differentiated to EpiLCs for 48 h in 35 mm dishes as described previously, in three biological replicates. Total RNA was extracted with Trizol and its integrity was evaluated by agarose gel electrophoresis. Samples were sent to Macrogen Inc for library construction and sequencing. Libraries were prepared with TruSeq stranded total RNA with Ribo-Zero Gold and sequenced on an Illumina NovaSeq6000 platform to deliver 30M 100 bp pair-end reads for each sample. Quality checked reads (FastQC) were processed in Fastp tool to remove any leftover adapters and low-quality ends. Processed reads were mapped to the mouse reference genome (GRCm39/mm39) using STAR aligner (Dobin et al., 2013) in two-pass mode, producing gene abundance quantification as well. Raw counts were then fed into DESeq2 for differential expression analysis (fdr<0.1). Graphics and statistical analyses were performed in R. Raw data will be made available upon publication of the manuscript. The list of DE genes is available in Table S1.

For the analysis of published ChIP-seq data, we used the ChIP-atlas web service (Zou et al., 2022). Bigwig and peak-called data for Oct6 EpiSCs binding profiles (SRX1410928) were downloaded from ChIP-atlas and analyzed in R using the dplyr and rtracklayer libraries. Peaks were assigned to nearby genes using GREAT (McLean et al., 2010). Crossing our RNA-seq database with Oct6 EpiSCs binding profiles determined that 114 out of 292 DEG contained Oct6 peaks. To ascertain the number of genes with Oct6 peaks out of 292 random genes we used a bootstrap strategy in R with 10.000 iterations (Efron and Tibshirani, 1993), which allowed us to determine the random distribution of Oct6 binding for that number of genes.

### Statistical analysis

Experimental results are presented as mean ± standard error of the mean (SEM) of at least three biological replicates or a representative experiment of the independent replicates. Statistical significance between groups was analyzed using either paired Student’s t-test or randomized block design (RBD) ANOVA. Residuals fitted normal distribution and homogeneity of variance. Comparisons between means were assessed using the Tukey test. Statistical analysis was performed using Infostat Software (JA di Rienzo et al., 2013).

### Tables

Table S1: DEG for WT and Oct6 KO EpiLCs

Table S2: Primers and sgRNA sequences

Table S3: Antibodies

Table S4: Nanog Stellaris Probes Quasar 670.

## Supplementary Figures

**Figure S1.**
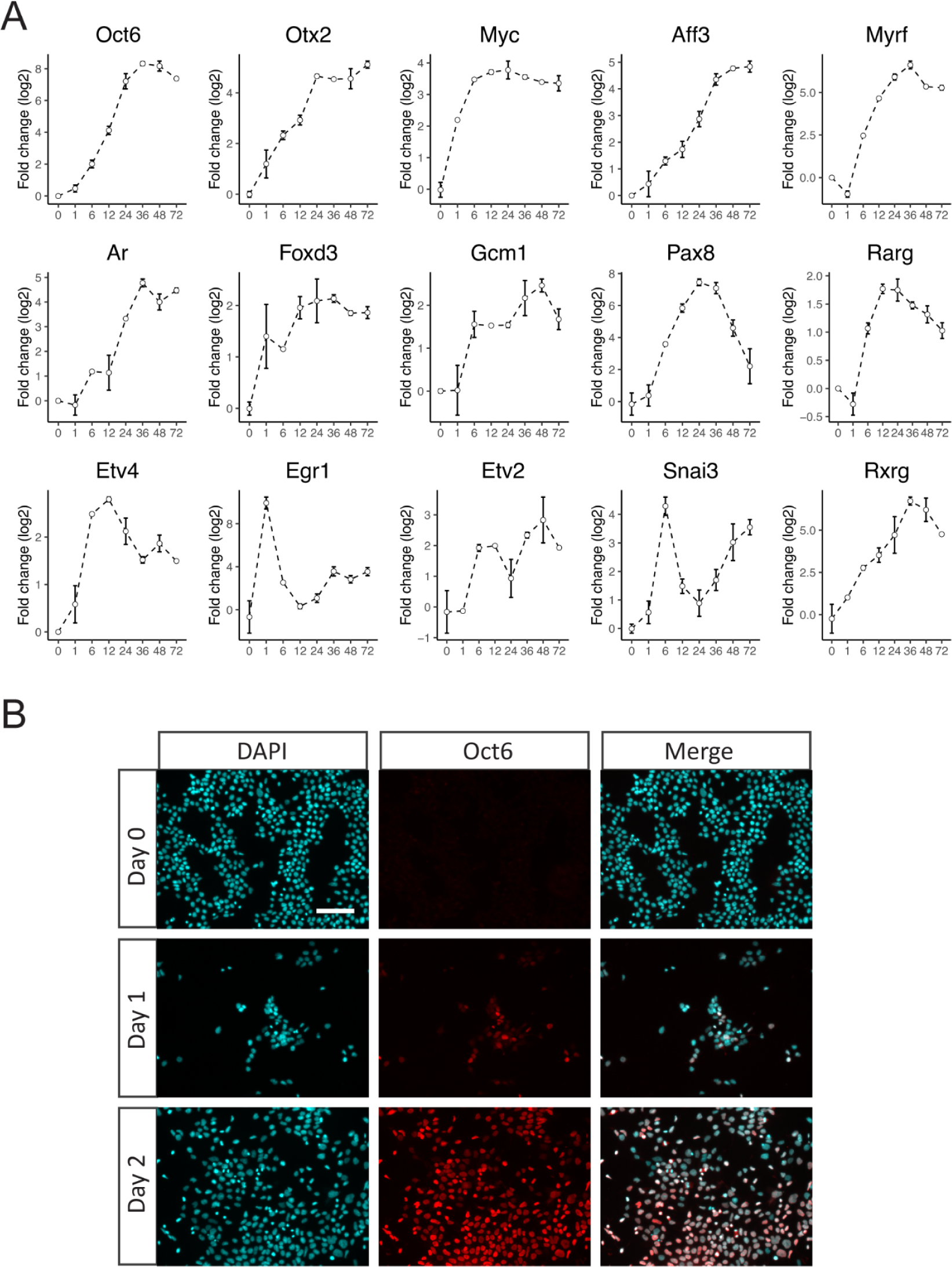
Expression of TFs induced during EpiLC differentiation. (A) Expression of TFs induced during the differentiation of EpiLCs. Re-analysis of RNA-seq data of Yang et al (Yang et al., 2019). (B) Representative immunofluorescence of OCT6 in undifferentiated cells and 24 or 48 h EpiLCs. Scale bar: 100 µm.

**Figure S2.**
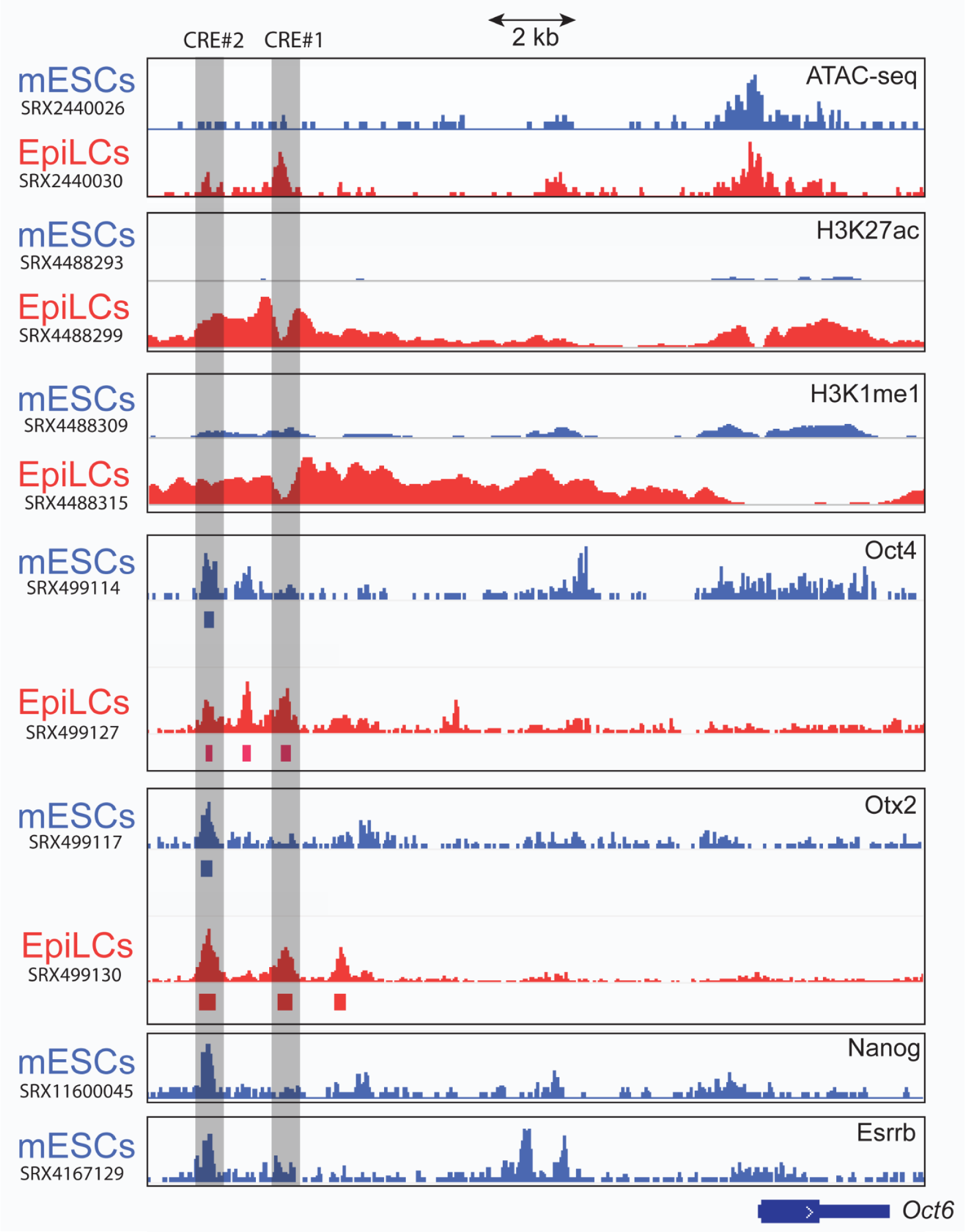
Binding of pluripotency TFs and histone marks at the Oct6 promoter. Evaluation of the binding of OCT4, OTX2, H3K27ac, H3K4me1, NANOG, and ESRRB at the Oct6 locus based on previously published ChIP-seq experiments (Atlasi et al., 2019; Bleckwehl et al., 2021; Buecker et al., 2014; Chen et al., 2018; Narita et al., 2021; Yang et al., 2019). Two CRE were identified. CRE#2 is bound by the four TFs in mESCs. OTX2 and OCT4 binding is extended to CRE#1 in EpiLCs, that also shows the topology of active enhancer marks H3K27ac and H3K4me1. The respective SRA experiment IDs are indicated on the left.

**Figure S3.**
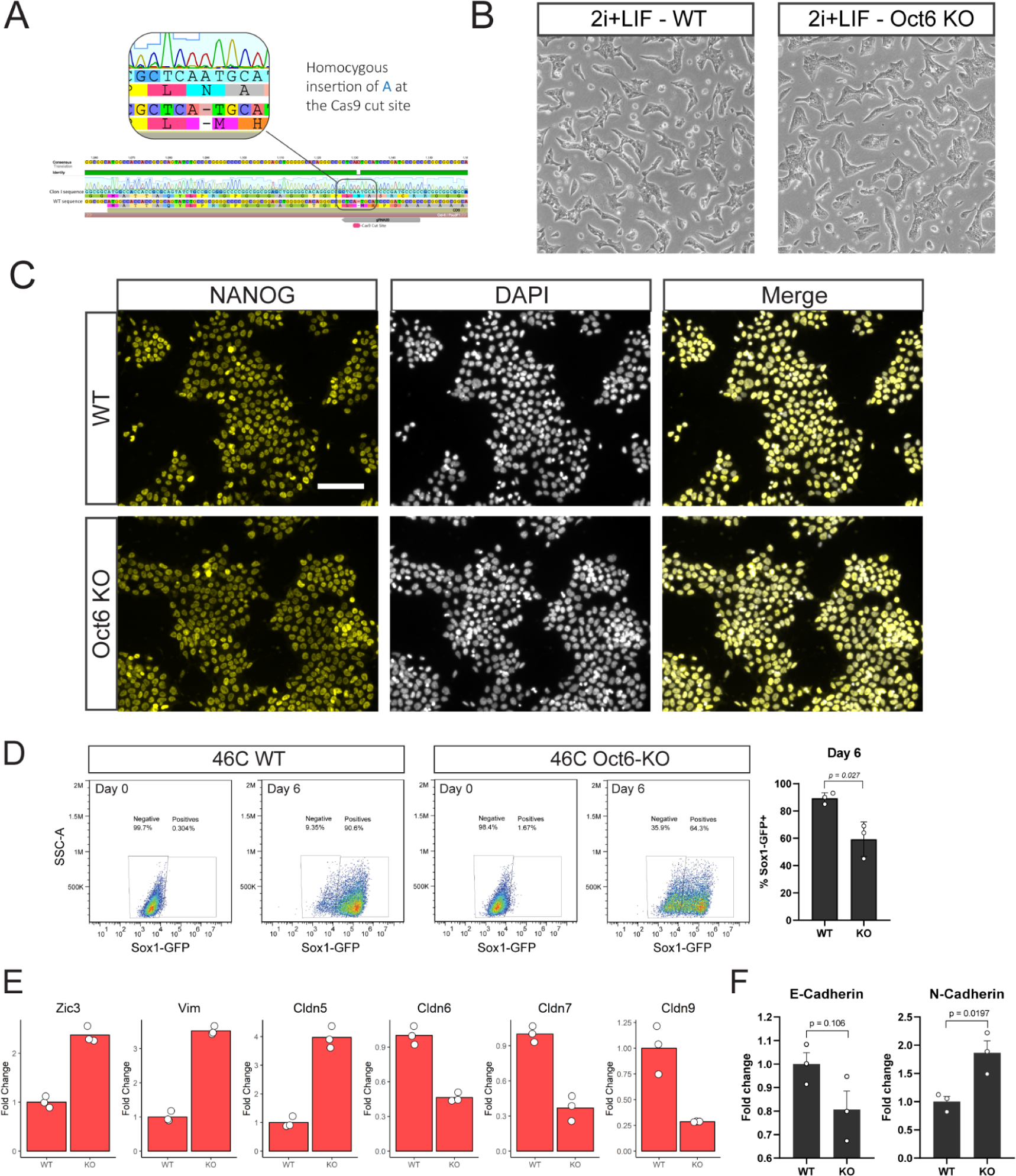
Generation and validation of Oct6 KO mESCs. (A) DNA sequencing of the Oct6 locus for WT 46C cells and for “Clon I” shows that the latter contains a homozygous insertion of an adenine at the Cas9 cutting site. This insertion generates a change in the open reading frame of Oct6 modifying the amino acid sequence starting from aminoacid 22 and generating a premature stop codon. (B) Brightfield images showing the similar morphology of WT and Oct6 KO mESCs cultured in 2i+LIF. (C) Immunofluorescence of the pluripotency markers OCT4, SOX2, NANOG, and KLF4 showing similar expression in WT vs Oct6 KO cells maintained in 2i+LIF. Scale bar: 100 µm. (D) Neural precursor differentiation was evaluated by flow cytometry analysis of the Sox1-GFP reporter. Comparison of WT and Oct6 KO 46C cells on day 0 (2i+LIF) or at day 6 of neural differentiation. *Left*, flow cytometry plots of a representative biological replicate. *Right*, percentage of Sox1-GFP+ neural precursors at day 6 for WT and Oct6 KO cells for three biological replicates. (E) RT-qPCR analysis of E-Cadherin and N-Cadherin in WT and Oct6 KO EpiLCs. Results are as presented as mean ± SEM for three independent replicates.

**Figure S4.**
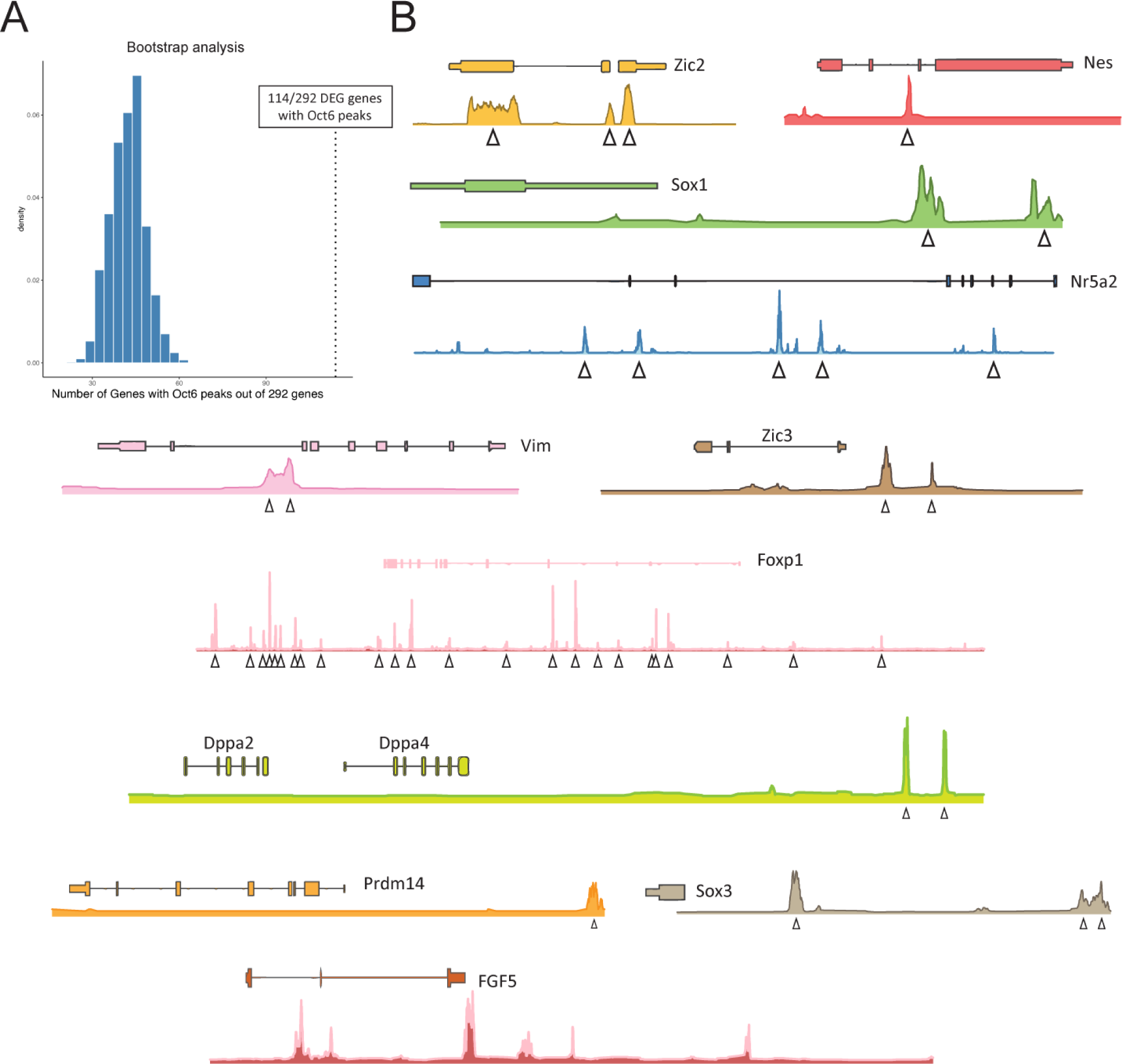
Binding of OCT6 to the genome in EpiSCs. (A) Analysis of DE genes of the RNA-seq data comparing WT vs Oct6 KO EpiLCs that contain OCT6 binding peaks in EpiSCs based on Matsuda et al. 114 out of 292 DE genes contained OCT6 binding peaks, suggesting a possible direct regulation by this TF. To obtain the random distribution of OCT6 binding among 292 randomly selected genes we followed a bootstrap strategy with 10.000 iterations. (B) OCT6 binding peaks in EpiSCs in example DE genes from our RNA-seq experiment.

**Figure S5.**
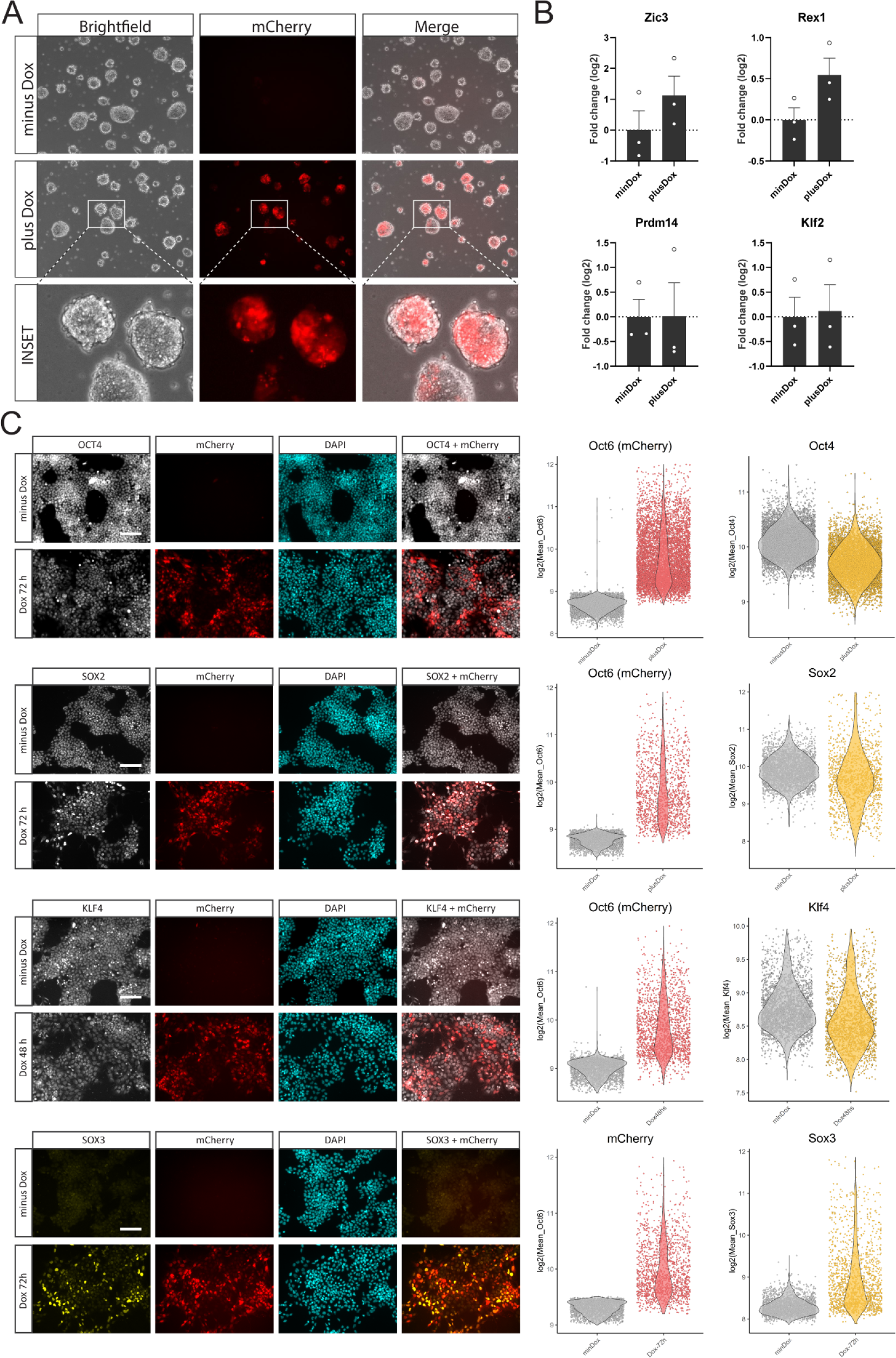
Overexpression of OCT6 in undifferentiated Oct6 KO cells. (A) Overexpression of the mCherry fluorescent protein (alone) after Dox treatment did not induce any morphological changes as cells presented the typical dome shape with no cell protrusions. (B) RT-qPCR analysis of Zic3, Rex1, Prdm14, and Klf2 in Oct6-P2A-mCherry cells maintained in 2i+LIF with or without Dox treatment. Results are as presented as mean ± SEM for three independent replicates. (C) Quantitative immunofluorescence of OCT4, SOX2, KLF4, and SOX3 in Oct6-P2A-mCherry cells maintained in 2i+LIF untreated or Dox treated for 24, 48, and 72 h. Panels on the left show representative immunofluorescence images. The right charts show the quantifications for mCherry and the selected TFs. Scale bar: 100 µm.

**Figure S6.**
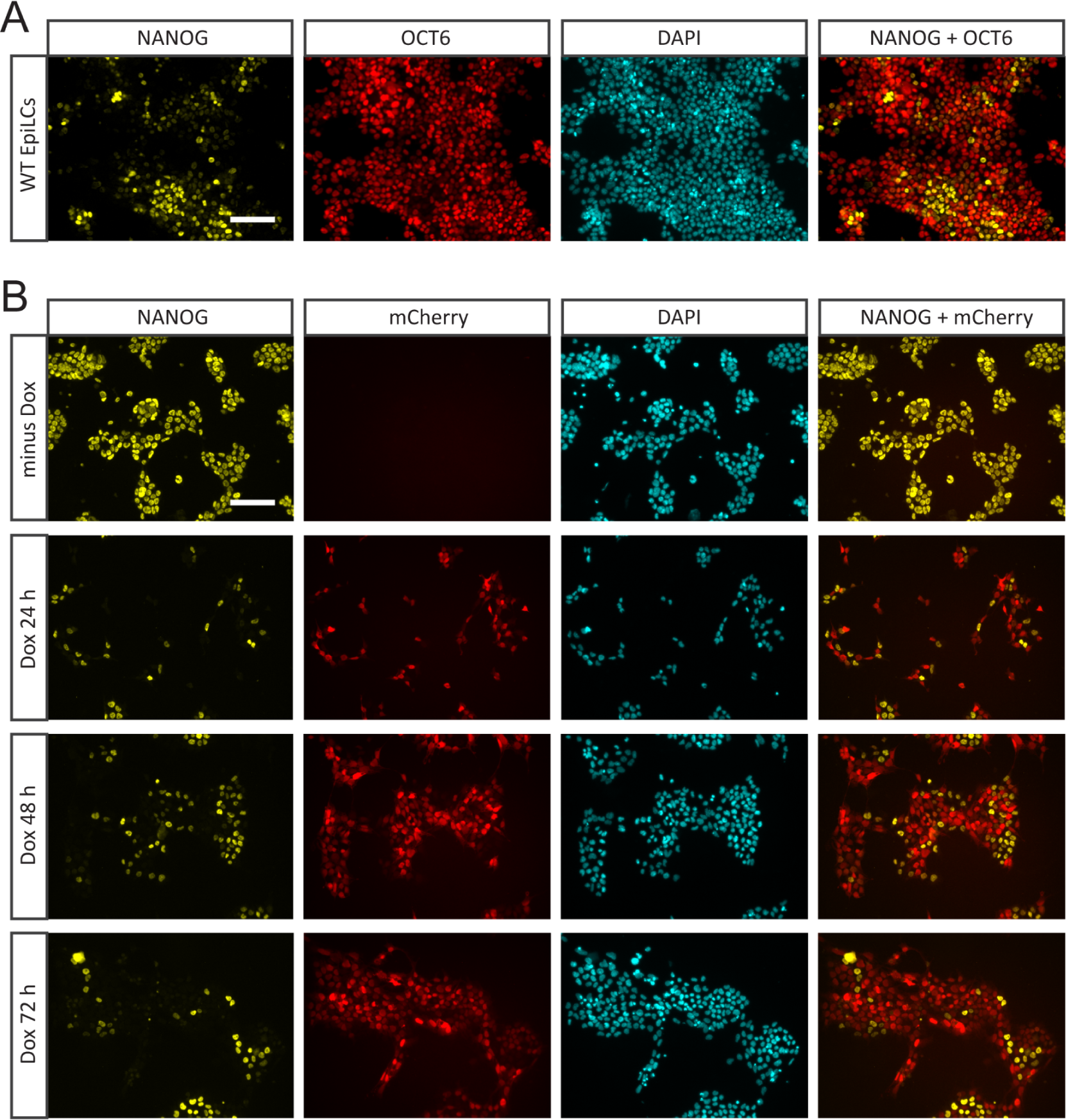
Oct6 and Nanog are expressed in a mutually exclusive manner. (A) Immunofluorescence analysis of OCT6 and NANOG in WT 48h EpiLCs showing mutually exclusive expression in normal conditions. Scale bar: 100 µm. (B) Inducible overexpression of Oct6-P2A-mCherry in Oct6 KO cells maintained in 2i+LIF shows that Oct6 represses the expression of Nanog. Cells were either untreated or treated with Dox for 24, 48, and 72 h. Scale bar: 100 µm.

**Figure S7.**
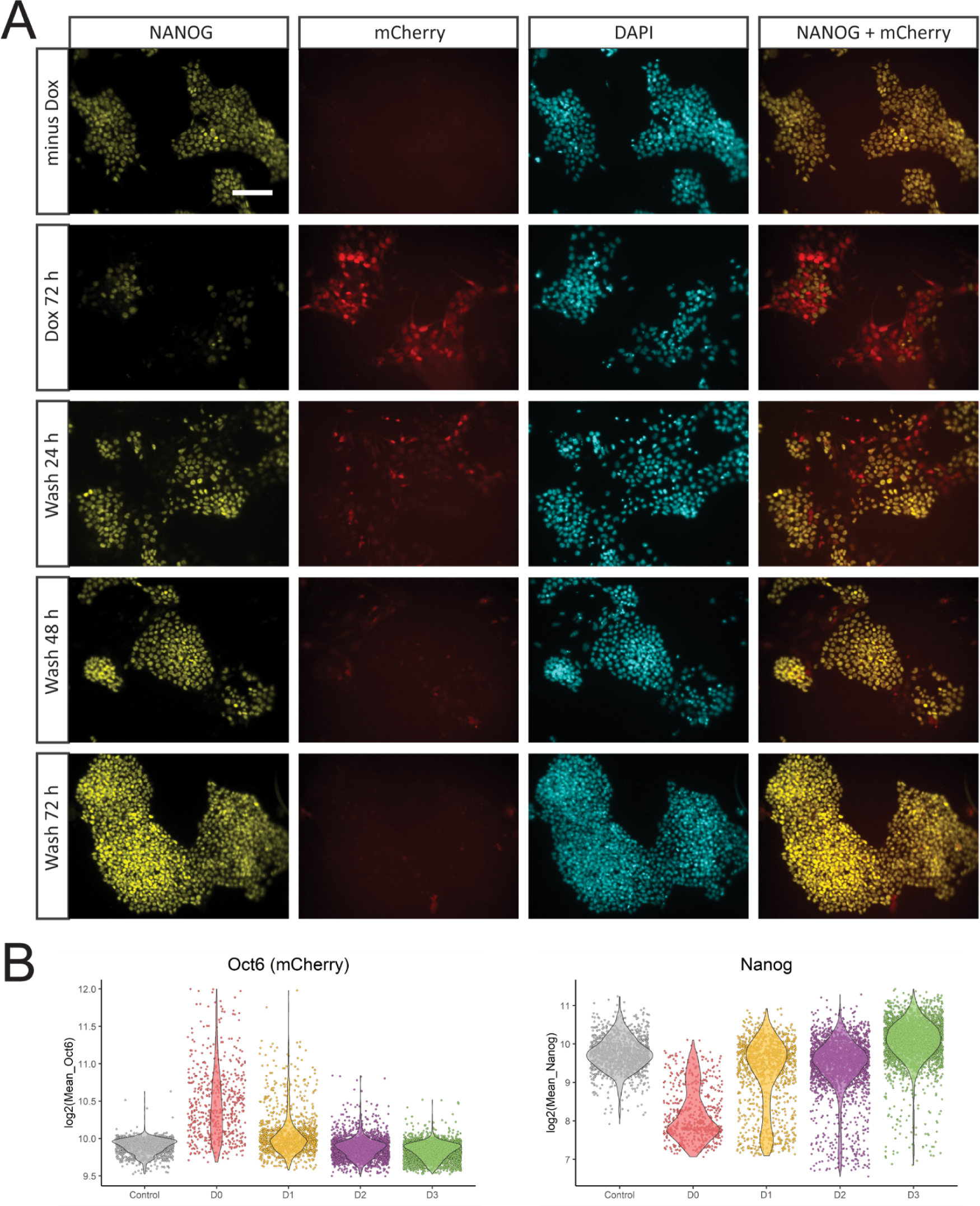
Repression of Nanog by Oct6 overexpression is reversible. (A) Oct6-P2A-mCherry cells maintained in 2i+LIF were untreated (minDox) or Dox treated for 72 hours and analyzed for mCherry and Nanog expression. Additionally, 72 h Dox treated cells were released from Dox induction and cultured for an additional 24, 48, or 72 h after washing and further evaluated for mCherry and Nanog expression. Representative images are shown. Scale bar: 100 µm. (B) Quantification of nuclear mCherry and NANOG signal of the experiment in A. mCherry fluorescence was rapidly reduced after Dox release. Nanog expression after Dox treatment is restored to similar levels as in untreated cells, showing that its repression by OCT6 is reversible.

**Figure S8.**
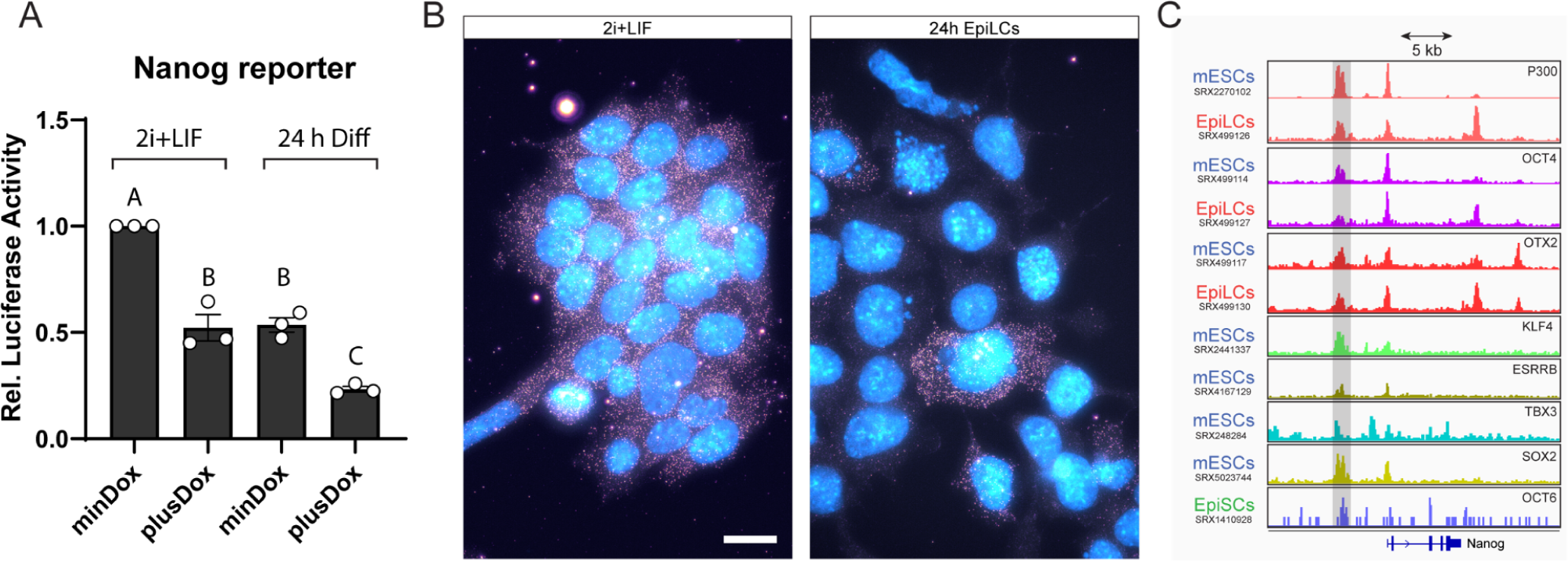
Single-molecule Nanog RNA fish and 3’UTR analysis of translational repression. (A) Analysis of the Nanog5P luciferase reporter. mESCs were co-transfected with Nanog5P-Luc and ePB-HA-Oct6-P2A-mCherry plasmids, maintained in either undifferentiated conditions or set to differentiate for 24 h, in the presence or absence of Dox. Firefly luminescence was assessed to evaluate Nanog promoter activity. The plot shows normalized luminescence levels relative to undifferentiated mESCs without Dox. Results are presented as mean ± SEM for three independent replicates. Significance between groups was analyzed by linear mixed models (LMM) and indicated with different letters. (B) Single-molecule RNA fish of Nanog of cells in 2i+LIF or after 24 h of EpiLCs induction. Individual Nanog mRNA transcripts can be observed in the cell’s cytoplasm. Transcript count is importantly reduced after 24 h of differentiation, with only a few cells maintaining Nanog expression. Nuclei were stained with DAPI (cyan). Scale bar: 20 µm. (C) Binding of different TFs at the Nanog locus based on previously published ChIP-seq experiments. The respective SRA experiment IDs are indicated on the left.

## Supplementary Tables

Table S1: DEGs RNA-seq

Table S2: Oligonucleotides

Table S3: antibodies

Table S4: Nanog smRNA FISH probes

